# Accurate estimation of SNP genotypes and genetic relatedness from DNA methylation data

**DOI:** 10.1101/2024.04.15.589670

**Authors:** Yi Jiang, Minghan Qu, Minghui Jiang, Xuan Jiang, Shane Fernandez, Tenielle Porter, Simon M. Laws, Colin L. Masters, Huan Guo, Shanshan Cheng, Chaolong Wang

**Affiliations:** Ministry of Education Key Laboratory of Environment and Health, School of Public Health, Tongji Medical College, Huazhong University of Science and Technology, Wuhan, China; Department of Epidemiology and Biostatistics, School of Public Health, Tongji Medical College, Huazhong University of Science and Technology, Wuhan, China; Centre for Precision Health, Edith Cowan University, Perth, WA, Australia; Collaborative Genomics and Translation Group, School of Medical and Health Sciences, Edith Cowan University, Perth, WA, Australia; Curtin Medical School, Bentley, WA, Australia; The Florey Institute of Neuroscience and Mental Health, University of Melbourne, Melbourne, VIC, Australia; Department of Occupational and Environmental Health, School of Public Health, Tongji Medical College, Huazhong University of Science and Technology, Wuhan, China

**Author notes:** These authors contributed equally to this work (Yi Jiang, Minghan Qu). Correspondence (C.W.), (S.C.).

**Keywords:** DNA methylation, genotype calling, genetic relatedness, population structure, epigenome-wide association study

## Abstract

Epigenome-wide association studies (EWAS) are susceptible to widespread confounding caused by population structure and genetic relatedness. Nevertheless, kinship estimation is challenging in EWAS without genotyping data. We propose MethylGenotyper, a method that for the first time enables accurate genotyping at thousands of SNPs directly from commercial DNA methylation microarrays. We model the intensities of methylation probes near SNPs with a mixture of three beta distributions corresponding to different genotypes and estimate parameters with an expectation-maximization algorithm. We conduct extensive simulations to demonstrate the performance of the method. When applying MethylGenotyper to Infinium EPIC array data of 4,662 Chinese, we obtain genotypes at 4,319 SNPs with a concordance rate of 98.26%, enabling the identification of 255 pairs of close relatedness. Furthermore, we show that MethylGenotyper allows for the estimation of both population structure and cryptic relatedness among 702 Australians of diverse ancestry. We have implemented MethylGenotyper in a publicly available R package to facilitate future large-scale EWAS.

## Introduction

DNA methylation (DNAm), which involves transferring a methyl group on to the C5 position of the cytosine, is an important mechanism of gene regulation and is dynamically variable in response to environmental changes. It has become the most widely studied type of epigenetic modifications owing to the development of high-throughput DNAm microarrays. Specifically, the Infinium HumanMethylation450 (450K) and HumanMethylationEPIC (EPIC) arrays can simultaneously assay DNAm levels at hundreds of thousands of Cytosine-phosphate-Guanine (CpG) sites across the genome. Using these arrays, epigenome-wide association studies (EWAS) have found numerous CpG sites in association with complex diseases or environmental exposures, leading to a better understanding of disease etiology [1-3]. Similar to genome-wide association studies (GWAS), population structure and cryptic relatedness can lead to spurious association signals in EWAS. DNAm levels might be divergent between populations due to the impacts from distinct genetic and environmental factors [4]. In contrast, related samples are likely to share similar environmental exposures and thus similar DNAm levels [1,5]. Despite the potential confounding effect, cryptic relatedness is often ignored in EWAS because there is no existing method to infer genetic relatedness based on DNAm data. Even in cohorts where both DNAm and GWAS data have been generated, the sample overlap between the two datasets is unlikely perfect due to their separate quality control (QC) procedures or other logistic factors. Therefore, methods to infer genetic relatedness directly from DNAm data will be very useful to facilitate large-scale EWAS.

With enough SNP genotypes, we can easily estimate genetic relatedness in samples with or without population structure using existing tools [6-10]. While the Infinium methylation arrays have been designed to incorporate SNP probes to facilitate the identification of sample swapping [11,12], the number of SNPs is too small to obtain accurate kinship estimates (*e*.*g*., EPIC array consists of 59 SNP probes, including 6 on the X chromosome). On the other hand, tens of thousands of CpGs near common SNPs (*i*.*e*., minor allele frequency MAF > 0.01) are often discarded by standard QC, because nearby SNPs can introduce mismatches to the probe sequence and thus interfere with the measured methylation intensity [13-15]. It has been found that the measured methylation intensities at these probes often show multi-modal distributions depending on the SNP genotypes, especially when common SNPs are present at the extension base [16-18]. Thus, we speculate that it is possible to infer SNP genotypes, and subsequently population structure and genetic relatedness, based on methylation intensity.

By design, there are two types of Infinium methylation probes. Type I probes use two beads at each locus to measure the methylated and unmethylated signals separately. Fluorescent colors are determined by the nucleotide at the extension base (*i*.*e*., red for A and T alleles, green for G and C alleles). Thus, SNPs (except for A/T and G/C SNPs) at the extension base will cause color channel switching (CCS). Genotypes for these SNPs can be accurately determined by comparing signal intensity from different color channels [13]. Type II probes use a single bead at each locus, with the extension base occurring at the target CpG. After bisulfite conversion, the red-labeled A and green-labeled G nucleotides will bind to the unmethylated and methylated alleles, respectively. In the presence of a SNP at the target CpG, a red color will be detected if the target C allele is mutated to A or T despite no effect of bisulfite conversion. On the other hand, a green color will remain detected if C is mutated to G. It is much more challenging to infer genotypes for Type II probes than for Type I probes, because both methylation and mutation can affect the fluorescent color of the single bead at Type II probes. To the best of our knowledge, there are no existing methods to estimate genotypes for SNPs near Type II probes, although the Infinium methylation arrays consist of many more Type II probes than Type I probes [14].

In this study, we develop a novel method named MethylGenotyper to perform genotype calling based on DNAm data for SNP probes, Type I probes, and Type II probes (**Figure 1**). For each type of probes, we first convert the methylation intensity signals to the Ratio of Alternative allele Intensity (RAI), which was first proposed by Zhou *et al*. [13] for Type I probes targeting CCS SNPs. RAI is expected to follow a three-modal distribution peaks near 0, 0.5, and 1, corresponding to three genotypes, respectively. We model RAI for each type of probes with a mixture of three beta distributions and one uniform distribution, and employ an expectation-maximization (EM) algorithm to obtain the maximum likelihood estimates (MLE) of model parameters and genotype probabilities. We evaluate the performance of the method in parameter estimation and genotype calling by extensive simulations with different sample sizes and numbers of SNPs. Next, we apply MethylGenotyper to two empirical datasets with both DNAm data (EPIC array) and GWAS data, namely the Dongfeng-Tongji (DFTJ) cohort [19], consisting of 4,662 Chinese samples, and the Australian Imaging, Biomarker & Lifestyle (AIBL) study [20,21], consisting of 702 samples with diverse ancestry. In both datasets, we demonstrate that MethylGenotyper can infer high-quality genotypes at over 4,000 SNPs, enabling accurate estimation of individual ancestry and pairwise relatedness. We have implemented MethylGenotyper into a publicly available R package to facilitate future large-scale EWAS.

**Figure 1.**
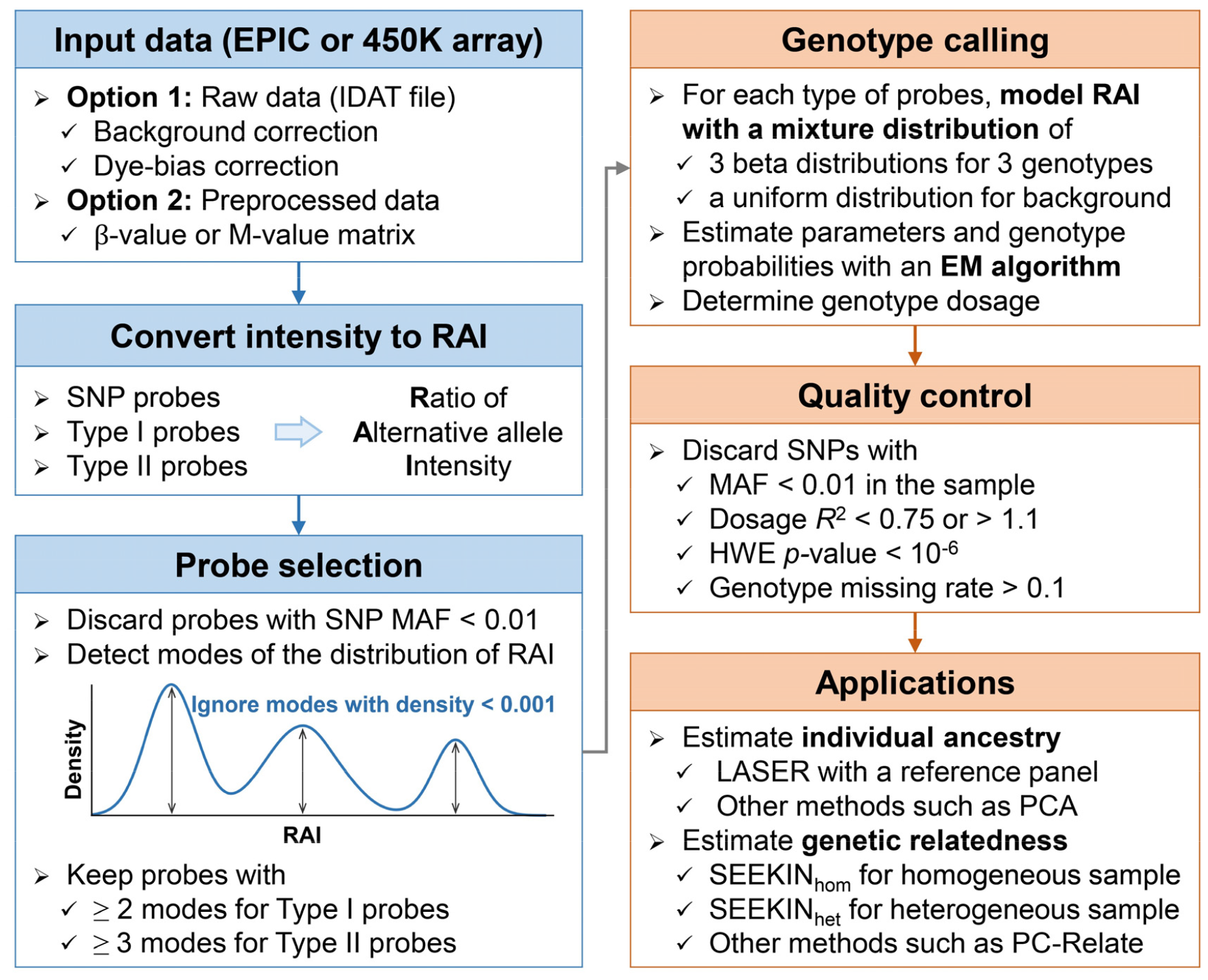
The workflow of MethylGenotyper. MethylGenotyper takes either raw intensity data (recommended) or preprocessed data (only works for SNP probes and Type II probes) to calculate the RAI of each probe. We keep probes with MAF > 0.01 in the corresponding population in 1KGP and require the RAI of each probe to have two or three modes. We develop an EM algorithm to fit RAIs of each type of probes with a mixture of three beta distributions and one uniform distribution. The three beta distributions correspond to three genotypes and their weights are probe-specific by assuming HWE. The uniform distribution represents background noise with a constant weight across the same type of probes.

## Results

### Correlation between DNAm intensity and SNP genotypes

We first examined squared correlation between the measured DNAm β-values and genotypes (coded as 0, 1, and 2) of nearby SNPs in the DFTJ dataset (**Figure S1**) [19]. We focused on biallelic autosomal SNPs with MAF > 0.01 in East Asian samples of the 1000 Genomes Project (1KGP), and excluded probes with multiple SNPs within 5 bp from the 3’ end of the probe. After regressing out age, sex, body-mass index (BMI), smoking status, sample plates and six immune cell type proportions, the highest *R*^2^ was observed for Type II probes with a SNP at the extension base (median *R*^2^ = 0.90 at position −1, **Figure S2A**), which was expected because Type II probes measured DNAm at the extension base. For Type I probes, the highest *R*^2^ was observed at the 3’ end (median *R*^2^ = 0.48 at position 0), where DNAm was measured. Importantly, the number of Type II probes with a SNP at the extension base (*n* = 7,619) is the largest among all probe categories we examined (**Figure S2B**). Considering both the *R*^2^ and the number of probes, we chose to focus on Type II probes with a SNP at the extension base, in addition to the SNP probes and Type I probes targeting CCS SNPs [13].

### Performance of MethylGenotyper in simulation data

We conducted two sets of simulations to examine MethylGenotyper based on the parameters of Type I probes and Type II probes obtained from real data (see next section), respectively. Our method fit the simulated RAI distributions perfectly for both probe types (**Figure S3**). The estimated values of ***α*** and ***β*** (parameters of three beta distributions), *λ* (relative intensity of the background noise), and ***ϕ*** (allele frequencies), approached their simulated true values as the sample size and the number of SNPs increased (**Figures 2** and **S4**). For Type II probes, error rates of the parameter estimates began to stabilize as the sample size reached 800 (**Figure 2A-G**). For a dataset consisting of 3,200 samples and 4,000 SNPs, the estimation errors relative to the true values were 0.0046 (95% confidence interval: 0.0036-0.0055) for ***α***, 0.0041 (0.0033-0.0049) for ***β***, 0.086 (0.084-0.088) for *λ*, and 0.031 (0.031-0.031) for ***ϕ***. The genotype concordance rates were ∼98.4% for different numbers of samples and SNPs (**Figure 2I**). For Type I probes, the genotype concordance rates were slightly lower (∼97.7%), possibly because of a higher level of background noise (*λ* = 0.025 for Type I probes versus 0.015 for Type II probes) and a smaller number of simulated SNPs (**Figure S4I**).

**Figure 2.**
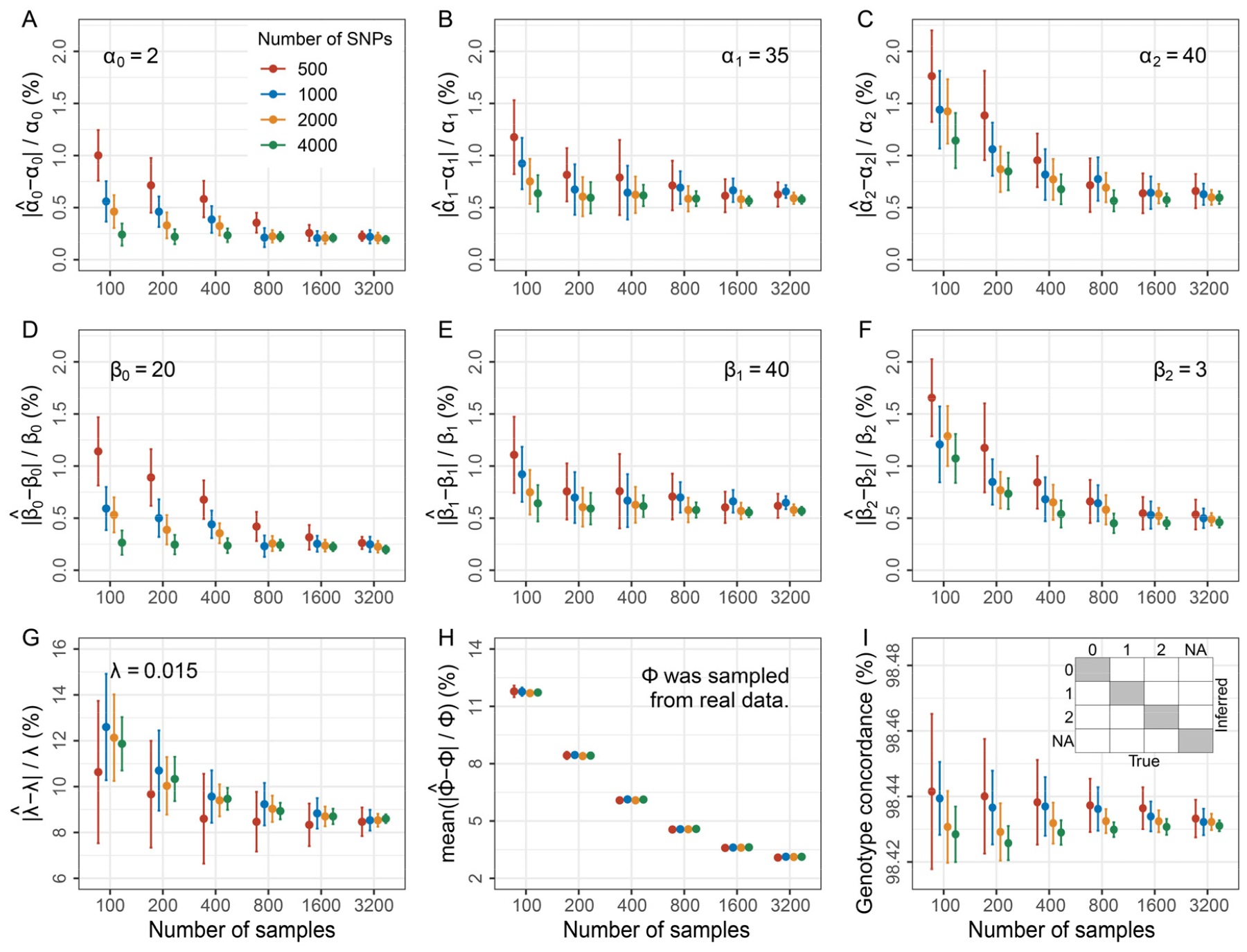
Performance of MethylGenotyper on simulation data mimicking the RAI distribution of Type II probes. We first simulated genotype matrices assuming HWE, with AFs (***ϕ***) randomly sampled from 1KGP. Conditional on the genotype matrices, RAI matrices were generated from a mixture distribution with parameters matching characteristics of Type II probes in the DFTJ dataset. **(A-G)** Error rates of the estimated parameters (***α, β***, *λ*). True parameter values were labelled in each panel. **(H)** Mean error rates of the estimated AFs. **(I)** Concordance of the inferred genotypes. As shown in the inserted panel, concordance was computed by dividing the number of genotypes in the shade areas by the total number of genotypes. In each panel, dots and vertical bars represent the means and 95% CIs (±1.96 standard errors), calculated from 20 repeats of simulation.

### Inferring genotypes using EPIC data from the DFTJ cohort

Following the probe selection procedure (**Figure 1** and **Materials and Methods**), we retained 53 SNP probes, 168 Type I probes, and 5,050 Type II probes for genotype calling using the EPIC data of 4,662 Chinese samples from the DFTJ cohort. The RAI distributions for each type of probes were fit well with parameter values shown in **Figure 3**. The relative intensity of the background noise *λ* was estimated to be 0.014, 0.024, and 0.017 for SNP probes, Type I probes, and Type II probes, respectively. After QC, we called genotypes at 4,319 SNPs, including 53 SNP probes, 111 Type I probes, and 4,155 Type II probes, with high accuracy (**Table 1**). Compared to the imputed GWAS data, the overall genotype concordance was 98.26% and the heterozygote concordance was 96.64%. We noted that the genotyping accuracy might be underestimated here, because erroneous genotypes were likely present in the imputed GWAS data. For Type II probes, the C allele at the extension base might be mutated to T, A or G alleles, of which C to T mutations accounted 93.3% of all SNPs. We found that SNPs with C to T or A mutations had similar genotype concordance rates at ∼98.27%, while C to G SNPs had lower genotype concordance at 97.75%. Given the high genotyping accuracy of MethylGenotyper, it was not surprised that the estimated allele frequencies were highly consistent with those based on the GWAS data for different probe types (**Figure 3D-F**).

**Table 1.** Accuracy of genotypes called by MethylGenotyper for the DFTJ dataset.

| **Probe type** | **Number of SNPs\*** | **Genotype concordance** | **Heterozygote concordance** |
| --- | --- | --- | --- |
| SNP probe | 53 (53) | 98.89% | 98.53% |
| Type I probe | 111 (109) | 98.26% | 96.93% |
| Type II probe | 4,155 (4,092) | 98.25% | 96.59% |
| C to T | 3,875 (3,818) | 98.27% | 96.62% |
| C to A | 193 (191) | 98.24% | 96.37% |
| C to G | 87 (83) | 97.75% | 96.05% |
| Overall | 4,319 (4,254) | 98.26% | 96.64% |

SNPs with $MAF < 0.01$ , HWE $p < 10^{-6}$ , $R^2 < 0.75$ or $> 1.1$ , or missing rate $> 0.1$ were excluded. Concordance was evaluated by comparison to the array genotyping data of the same samples.
\* Number of SNPs with array genotyping data were shown in the parentheses.

**Figure 3.**
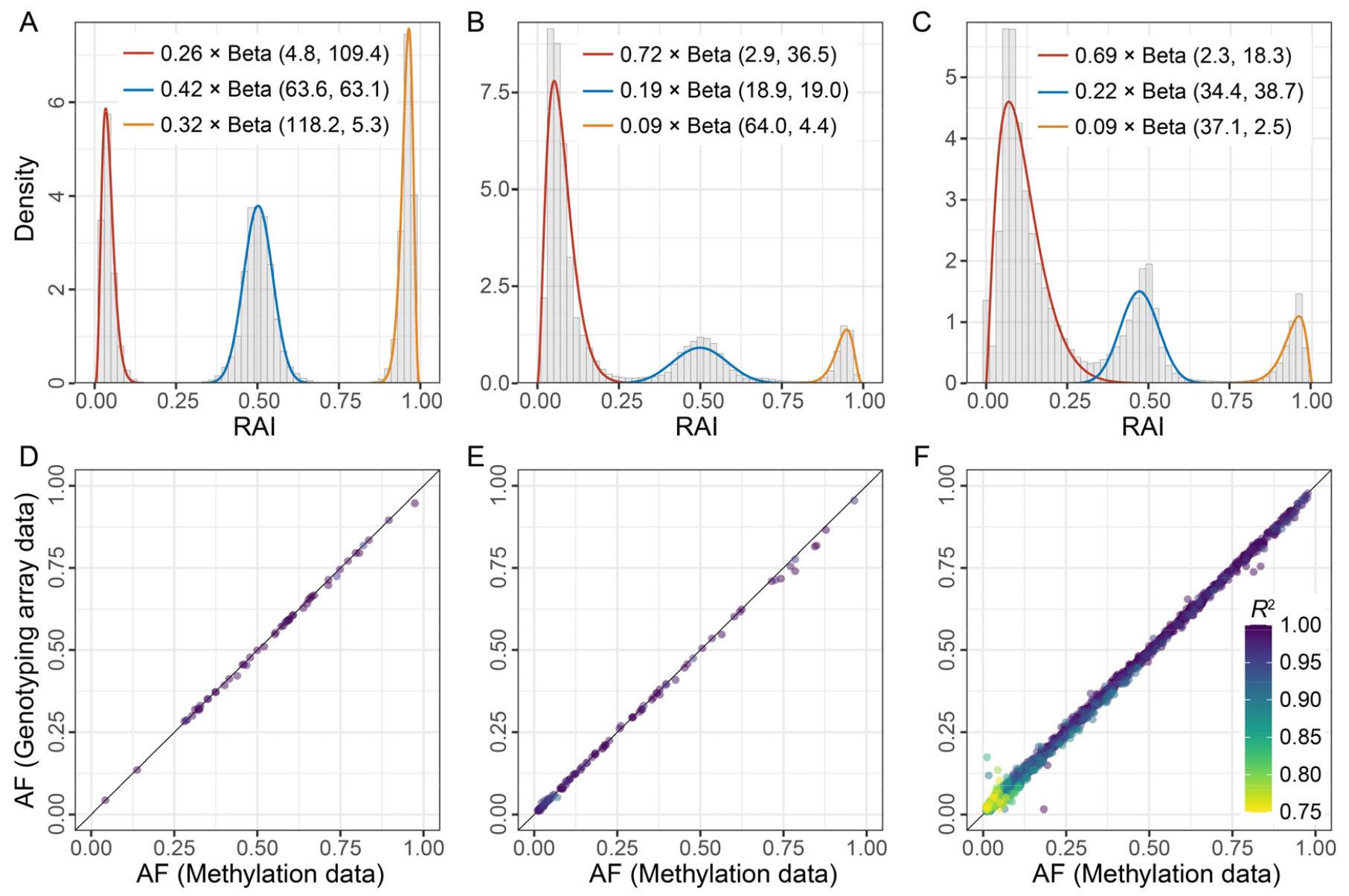
Performance of MethylGenotyper in the DFTJ dataset. **(A-C)** Fitted distributions of RAI for SNP probes **(A)**, Type I probes **(B)**, and Type II probes **(C)**, respectively. Histograms show the distributions of RAI for all selected probes and smooth lines indicate the fitted beta distributions, with weights averaged across probes. **(D-F)** Comparison of AFs derived from MethylGenotyper with those directly from array genotyping data for SNP probes **(D)**, Type I probes **(E)**, and Type II probes **(F)**, respectively. Each point represents a SNP, colored by the estimated dosage *R*^2^. For the bottom panels, only SNPs passed QC were shown.

### Estimation of genetic relatedness in the DFTJ dataset

Based on GWAS data, we identified 123 duplicated pairs, 110 pairs of 1^st^ degree, 22 pairs of 2^nd^ degree, and 53 pairs of 3^rd^ degree relatedness among 4,662 DNAm samples in the DFTJ cohort (**Figure 4A**). In contrast, based on genotypes inferred from 53 SNP probes and 111 Type I probes, it was difficult to distinguish related pairs from the huge number of unrelated pairs, with almost zero precision to identify 2^nd^ degree or closer relatedness (**Figure 4B-C, Figure S5A-B**). By additionally including the genotypes inferred from 4,155 Type II probes, the variance of kinship estimates was dramatically reduced, leading to clear separation of different degrees of relatedness (**Figure 4D, Figure S5C**). Compared to the benchmark using GWAS data, we could identify 2^nd^ degree or closer relatedness with 0.9659 precision and perfect recall rate (*F*_1_ = 0.9827).

**Figure 4.**
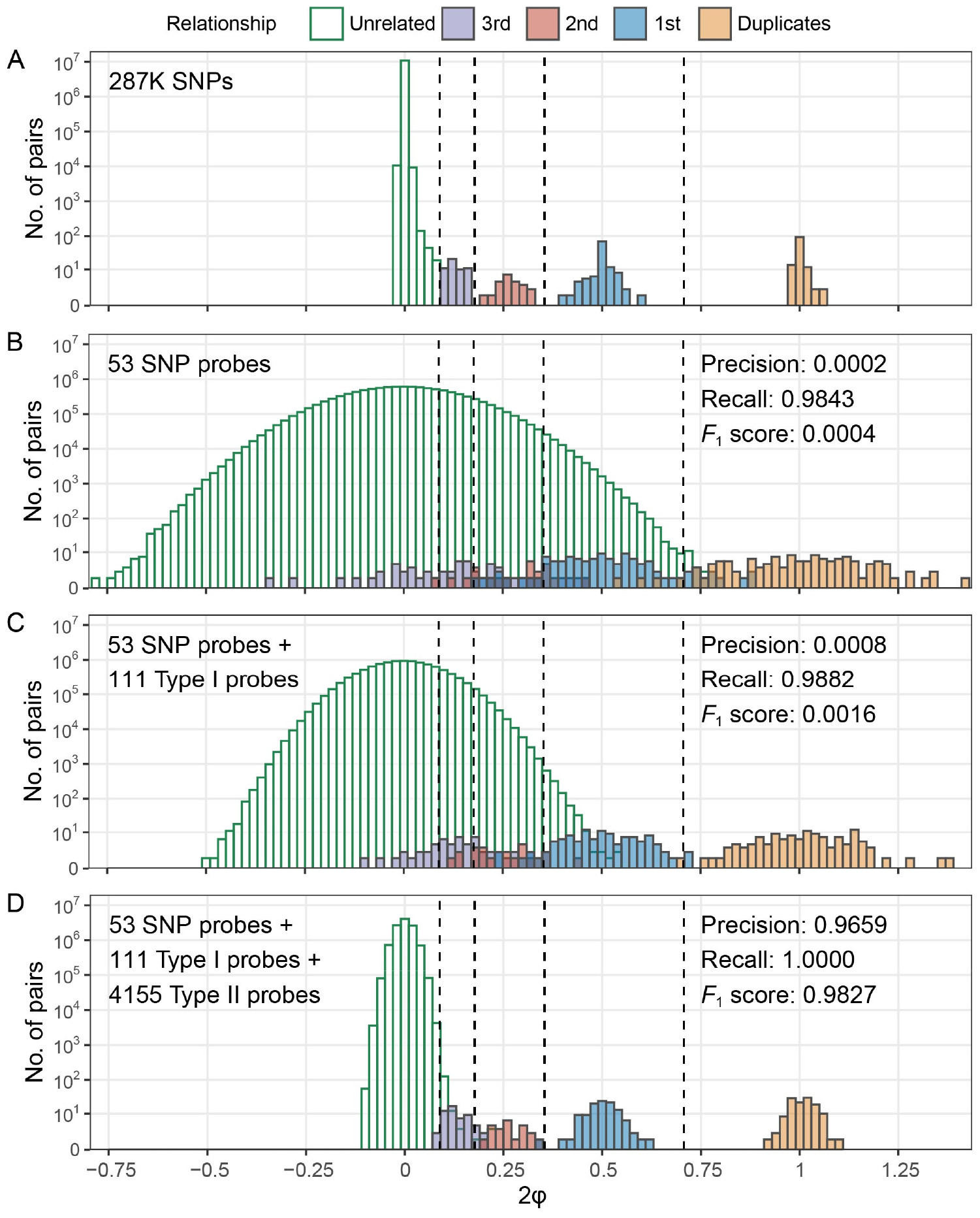
Estimation of genetic relatedness among samples in the DFTJ dataset. **(A)** Kinship estimates based on 286,727 genome-wide SNPs (gold standard). **(B)** Kinship estimates based on 53 SNP probes. **(C)** Kinship estimates based on 53 SNP probes and 111 Type I probes. **(D)** Kinship estimates based on 53 SNP probes, 111 Type I probes, and 4,155 Type II probes. Relationship types were determined by the gold standard kinship estimates in **(A)**, with the inference criteria indicated by vertical dashed lines. In (**B-D)**, precision, recall, and *F*_1_ score were calculated by comparison to the gold standard, treating 2^nd^ degree or closer relatedness as positive.

We also assessed whether our method could allow for accurate kinship estimation using the Infinium 450K array. By design, over 90% of the probes on the 450K array were included on the EPIC array [14]. Thus, we extracted DNAm data of these probes from the EPIC data of the DFTJ cohort and applied MethylGenotyper to call genotypes. We identified 2,212 SNPs with high-quality genotypes, including 53 from SNP probes, 104 Type I probes, and 2,055 Type II probes. The performance in kinship estimation based on these SNPs was similar to that based on the full EPIC data despite of slightly larger variance (**Figure S6**).

### Inference of population structure and cryptic relatedness in the AIBL dataset

We further validated MethylGenotyper using data from the AIBL study [22], in which both DNAm data (EPIC array) and GWAS data were available for 702 Australian samples. Based on the DNAm data, we obtained high-quality SNP genotypes at 4,217 probes, including 52 SNP probes, 135 Type I probes, and 4,030 Type II probes. The allele frequencies estimated from DNAm data were highly consistent with those from 1KGP European data, except for a small number of SNPs (**Figure S7**).

We first investigated the ancestral background of the AIBL samples using the LASER method with 1KGP data as the reference panel [23]. Surprisingly, while we found most samples clustered with Europeans, a handful of the samples clustered with East Asians, South Asians, or in between (**Figure 5A**). To account for the diverse ancestry background, we estimated kinship coefficients among AIBL samples using estimators for heterogeneous samples [6,10]. We found the kinship estimates were highly consistent between those derived from DNAm data and from GWAS data (**Figure 5B**). We identified four pairs of 1^st^ degree relatedness that were previously unknown. These results demonstrated the robustness of MethylGenotyper and its potential applications in the inference of population structure and cryptic relatedness among samples from diverse populations.

**Figure 5.**
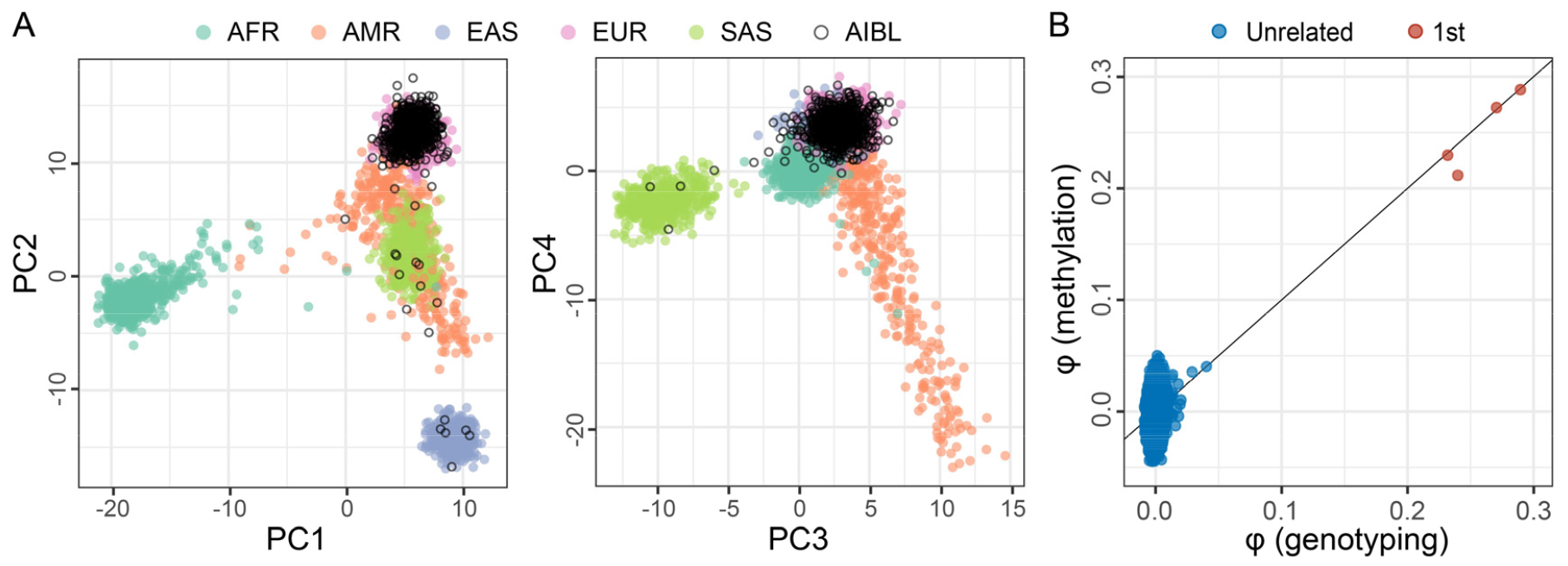
Performance of MethylGenotyper in the AIBL dataset. **(A)** Inferred ancestry of 702 AIBL samples in the ancestral space generated by the top four PCs of the 1KGP samples. Analysis was based on 4,217 SNPs called by MethylGenotyper. Abbreviations: AFR, Africans; AMR, Americans; EAS, East Asians; EUR, Europeans; SAS, South Asians. **(B)** Comparison of kinship estimates based on genotypes called by MethylGenotyper and those from genome-wide array genotyping data. Relationship types were determined based on array genotyping data.

## Discussion

Population structure and cryptic relatedness are major confounding factors in both GWAS and EWAS, where hundreds of thousands of sites are tested for phenotype association [1]. But genetic data are often not available or incomplete in EWAS samples. Several studies have explored the potential to infer population structure directly from DNAm data, mostly based on PCA of CpG sites near known SNPs [24-26]. The methods developed in these studies, without properly modelling the relationship between SNP genotypes and DNAm signals, have limited resolution to infer population structure because DNAm intensity can be affected by many other factors, including batch effects. Furthermore, no study has attempted to infer genetic relatedness directly from DNAm data, even though close genetic relatedness often implies shared environmental exposures that can affect both DNAm and phenotypes. In this study, we develop a novel method called MethylGenotyper to accurately infer genotypes at thousands of SNPs, based on DNAm data that are often discarded by standard QC. We demonstrate that SNP genotypes inferred by our method allow for accurate inferences of both population structure and genetic relatedness, and thus addressing a major confounding issue in EWAS.

While it has been noted that SNPs near CpG target sites can interfere methylation intensity [16-18], few studies have explored genotype calling at these SNPs, except for two studies [12,13]. Heiss and Just [12] developed ewastools to call genotypes specifically for tens of SNP probes incorporated into the Illumina 450K and EPIC arrays, and showed these SNPs can be used to identify mislabeled or contaminated samples. Zhou *et al*. [13] proposed to infer genotypes at hundreds of SNPs that can cause CCS at Type I probes. Nevertheless, based on EPIC array data, we showed that SNP probes and Type I CCS probes together were not sufficient for accurate kinship estimation to separate closely related and unrelated pairs. In contrast, we expand the number of genotyped SNPs by 25 times by incorporating thousands of Type II probes.

We process SNP probes, Type I probes, and Type II probes separately but under a unified statistical framework. We first generalize the RAI statistic proposed by Zhou *et al*. [13] for Type I probes to all three types of probes. We then model RAI for each type of probes with a mixture of three beta distributions and one uniform distribution, similar to the model in ewastools [12], except that we introduced probe-specific weights based on allele frequencies from external source to improve genotyping accuracy. With a sophisticated model and an EM algorithm, our method can infer genotypes with over 98% concordance rate for over 4,000 SNPs from EPIC array, allowing for almost perfect identification of *≤* 2^nd^ degree relatedness in the DFTJ dataset. Notably, similar performance in kinship estimation can be achieved even when we use DNAm probes available on the Illumina 450K array, supporting wide applicability of MethylGenotyper to different methylation arrays.

Based on EPIC data from the AIBL cohort, we further illustrated that SNP genotypes inferred by MethylGenotyper can be used to infer population structure and close relatedness among samples with diverse ancestry. We used the LASER method [23] to estimate individual ancestry in a reference ancestral space of worldwide populations, and the SEEKIN-het estimator [6] for kinship estimation, accounting for individual-specific ancestry background. The analysis workflow incorporating LASER and SEEKIN methods has been implemented in the MethylGenotyper package to facilitate the research community. Furthermore, while unexplored in the present study, we expect that high-quality genotypes from over 4,000 SNPs will be sufficient to identify fine-scale population structure within continental groups based on down-sampling experiments in a previous study [27].

In addition to estimating population structure and genetic relatedness, the accurate genotypes called by MethylGenotyper can be used in many downstream analyses, including estimation of inbreeding coefficient, and detection of sample contamination or sample swapping. Nevertheless, the utility of these genotypes in methylation quantitative trait locus (meQTL) mapping is limited due to the relatively small number of SNPs and the potential impacts of these SNPs on the methylation measurements of nearby CpGs.

The computational cost of our method increases linearly with the number of samples (*n*) and candidate probes (*m*), resulting in a complexity of O(*mn*). Using EPIC data of 1,000 samples as an example, the raw data preprocessing (background and dye bias correction) takes ∼17 minutes with 10 CPUs, while genotype calling takes ∼13 minutes with 1 CPU. The test is run on a high-performance computing cluster with Intel Xeon CPUs (2.3 GHz).

In conclusion, we have developed MethylGenotyper to accurately infer genotypes at thousands of SNPs directly from DNAm microarray data. We demonstrate that these SNP genotypes can be used to accurately estimate population structure and genetic relatedness, beyond simple tasks such as identifying mislabeled or contaminated samples. One limitation of MethylGenotyper is that we only focus on SNPs at the extension base of both Type I and Type II probes. Future studies might consider incorporating SNPs present at other nearby positions, which are also known to interfere with the measured DNAm intensity. Given the widespread confounding effects caused by population structure and genetic relatedness, we recommend the research community to incorporate MethylGenotyper into the standard analysis pipeline of EWAS to maximize statistical power and avoid spurious association signals.

## Materials and Methods

### DNAm and GWAS data from the DFTJ cohort

The DFTJ cohort is a prospective cohort of retired workers from the Dongfeng Motor Corporation in Shiyan, Hubei, China [19]. A total of 38,295 participants were enrolled in 2013, and leukocytes from 5,200 samples were selected for DNAm profiling using the Infinium HumanMethylationEPIC BeadChips in two batches (Fig. S1). We excluded samples with low detection quality (detection *P* > 0.01 in over 1% probes), mismatched methylation-inferred and self-reported sexes, discordant genotypes at >10 SNP probes compared with GWAS data, and those with no GWAS data, resulting in 4,662 samples. These 4,662 samples correspond to 4,542 unique individuals, including 114 with duplicate measurements and three with triplicate measurements. The raw IDAT files were processed by background correction with the *noob* method [28,29] and dye bias correction with the *RELIC* method [30]. Probes on the sex chromosomes were excluded.

GWAS data of 33,114 samples in the DFTJ cohort were assayed using the Infinium OmniZhonghua-8 arrays. After excluding low-quality samples (> 0.05 discordance rate at duplicate sites, call rate < 0.9, inbreeding coefficient < −0.1 or > 0.3 based on autosomal SNPs or < −0.2 based on the X chromosome), duplicated samples, and sex-mismatched samples, a total of 31,155 individuals passed QC. We excluded SNPs with call rate < 0.95, minor allele count (MAC) < 3, or Hardy-Weinberg Equilibrium (HWE) *P* < 10^−6^, leaving 775,059 autosomal SNPs and 24,134 X-chromosomal SNPs. We then phased and imputed the GWAS data by Eagle (v2.4.1) [31] and Minimac4 [32], with a whole-genome sequenced reference panel consisting of 3,931 East Asian samples from 1KGP [33] and the SG10K Project [34]. After excluding 20,640 variants with > 0.2 difference in allele frequency compared to East Asians in 1KGP, 14,495,888 variants with imputation *R*^2^ *≥* 0.3 and MAF *≥* 0.001 were kept for downstream analyses.

### DNAm and GWAS data from the AIBL study

The AIBL study (https://aibl.org.au) is a consortium between Austin Health, CSIRO, Edith Cowan University, the Florey Institute (The University of Melbourne), and the National Ageing Research Institute in Australia, aiming to improve the understanding of Alzheimer’s Disease [20,21]. We downloaded DNAm array data (EPIC array) of 726 samples from the Gene Expression Omnibus (GEO) repository (accession number: GSE153712) [22]. We downloaded IDAT files and processed the data with the *noob* method [28,29] and the *RELIC* method [30] for background correction and dye bias correction, respectively. A total of 716 samples in the AIBL cohort were genotyped using the Infinium OmniExpressExome arrays [35,36], including 702 samples that overlapped with the samples having DNAm data.

### Association between DNAm intensity and nearby SNPs

We examined the association between DNAm β-values and the genotypes of nearby SNPs based on the DFTJ data. We focused on biallelic SNPs with MAF > 0.01 in the East Asian samples of 1KGP [33] and excluded probes with multiple SNPs within 5 bp from the 3’ end of the probe. For each probe, we pre-adjusted β-values by regressing out sex, age, BMI, smoking status, sample plates, and six immune cell type proportions estimated by Bigmelon [37]. We computed the squared Pearson correlation (*R*^2^) between the residualised β-values and the genotype dosages of the nearby SNP as a function of the SNP position relative to the probe.

### Calculation of RAI

We consider three types of probes for SNP genotyping: (1) SNP probes by design, (2) Type I probes targeting CCS SNPs introduced by Zhou *et al*. [13], and (3) Type II probes with a SNP at the extension base. The Ratio of Alternative allele Intensity (RAI) was defined for each type of probes separately.

For SNP probes, the reference and alternative alleles are targeted by different probes and the RAI can be calculated as

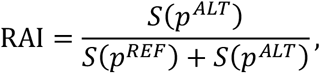

where *S*(*p*^*REF*^) and *S*(*p*^*ALT*^) denote probe signals supporting the reference allele and the alternative allele, respectively.

For Type I probes with CCS SNPs, we follow Zhou *et al*. [13] to calculate RAI as

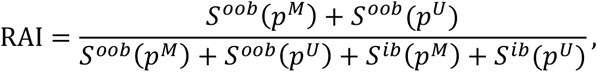

where *p*^*M*^and *p*^*U*^denote methylated and unmethylated probes, respectively, *S*^*oob*^is the out-of-band signal supporting the alternative allele, and *S*^i^ is the in-band signal supporting the reference allele.

For Type II probes, the extension base targets the cytosine of a CpG. Without mutation at the target site, a red color signal will be detected after bisulfite treatment when there is no methylation, while a green color signal will be detected when the cytosine is methylated. When C is mutated to A or T, a red color signal will always be detected. When C is mutated to G, a green color signal will always be detected. Thus, we have

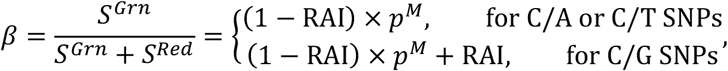

where *S*^*Red*^ and *S*^*Grn*^ represent the red and green color intensities, respectively, *p*^*M*^ represents the methylated proportion of C alleles, and *β* is the standard beta value calculated as the proportion of green color intensity. RAI roughly corresponds to 0, 0.5, and 1 for reference homozygotes, heterozygotes, and alternative homozygotes, respectively. Therefore, we expect *β* to follow a three-modal distribution with modes near (0, 0.5*p*^*M*^, and *p*^*M*^) for C/A or C/T SNPs and (*p*^*M*^, 0.5*p*^*M*^ +0.5, and 1) for C/G SNPs, assuming *p*^*M*^ is stable across samples for each CpG. In practice, we use the “multimode” method to check the distribution of *β* values for each CpG, including the number of modes, mode locations, and the density of each mode [38]. We keep probes that have at least two modes with density height > 0.001 (bandwidth = 0.04). For probes with more than three modes detected, we remove the lowest modes until only three modes remained. We define *l*_het_ as the location of the central mode of *β* (or the mode closest to 0.5 if only two modes are detected), which corresponds to heterozygotes. We then propose to estimate *p*^*M*^ for each CpG as

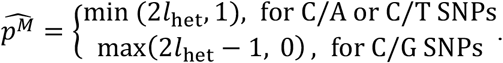

Finally, we calculate RAI as

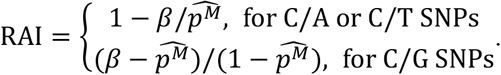

We truncate RAI values at 0.01 or 0.99 for those outside the range between 0.01 and 0.99.

For both Type I and Type II probes, we applied an additional filtering step based on the distribution of RAI values. We retained only Type I probes with at least two modes and Type II probes with at least three modes. We use a more stringent threshold for Type II probes because the SNPs for which we call genotypes correspond directly to the methylation target sites, which are more susceptible to the influence of methylation β values and potential confounding.

### Modelling the distribution of RAI

For each type of probes, we code the RAI values by an *m*×*n* matrix ***X***, where *m* and *n* indicate the number of probes and samples, respectively. We assume ***X*** follows a mixture of three beta distributions, corresponding to three genotypes, and a uniform distribution representing the background noise [12]:

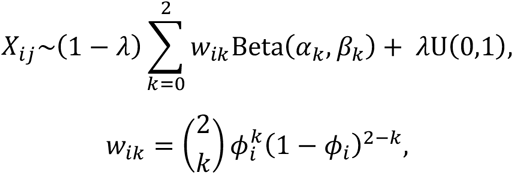

where *X*_*ij*_ represents the RAI value at probe *i* of sample *j, k* represents the genotype dosage, coded as 0, 1, and 2 for reference homozygotes, heterozygotes, and alternative homozygotes, respectively, Beta(*α*_*k*_, *β*_*k*_) represents the beta distribution with parameters *α*_*k*_ and *β*_*k*_, U(0,1) represents the standard uniform distribution, *λ* represents the probability that RAI comes from background noise, (1 - *λ*) *w*_*ik*_ represents the probability that RAI comes from genotype *k* with weight *w*_*ik*_ specified by the Hardy-Weinberg proportions at probe *i*, and *ϕ*_*i*_ represents the allele frequency at probe *i*.

### The EM algorithm

Let *B*_*ijk*_ = Beta(*X*_*ij*_; *α*_*k*_, *β*_*k*_) denote the probability density of Beta(*α*_*k*_, *β*_*k*_) at *X*_*ij*_. Assuming *X*_*ij*_ are independent, the log-likelihood function can be written as

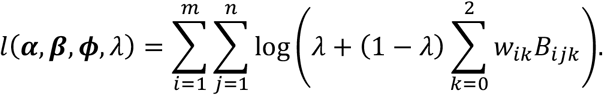

We develop an EM algorithm to estimate the parameters (***α, β, ϕ***, *λ*), of which the initial values were set as *α*_O_ = 5, *β*_O_ = 60, *α*_1_ = *β*_1_ = 30, *α*_2_ = 60, *β*_2_ = 5, *ϕ*_*i*_ = 0.2 for any *i*, and λ = 0.01.

In the E-step, we calculate the probability of *X*_*ij*_ from U(0,1) as

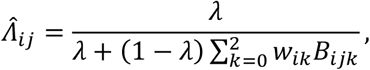

and the probability of *X*_*ij*_ from Beta(*α*_*k*_, *β*_*k*_) as

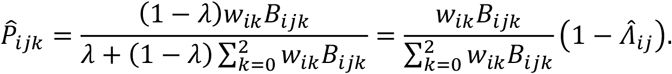

In the M-step, we update the parameters with their moment estimators:

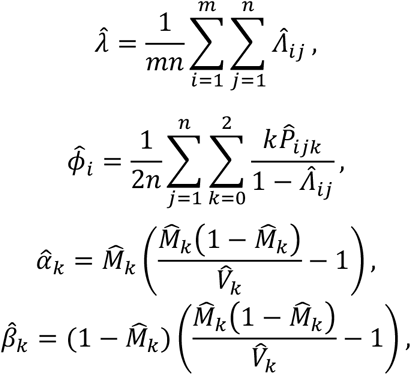

where

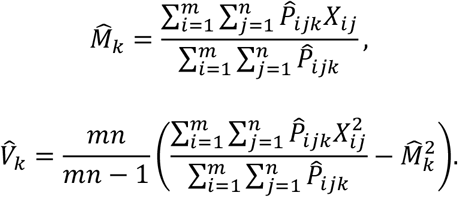

We iterate the E-step and M-step until the log-likelihood converges to the maximum, and thus obtain the maximum likelihood estimates of (***α, β, ϕ***, *λ*).

### Genotype calling and quality controls

With the estimated genotype probabilities 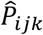 and background probabilities 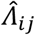, we set the genotypes with 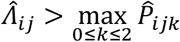 as missing and then update each 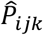 by dividing it with 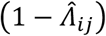 to make sure 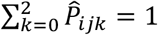 for any probe *i* and sample *j*. For the other non-missing genotypes, we denote the 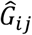 as the most probable genotype and 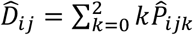 as the genotype dosage. Following Li *et al*. [39], we compute the allele frequency 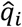 and the dosage 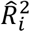 as

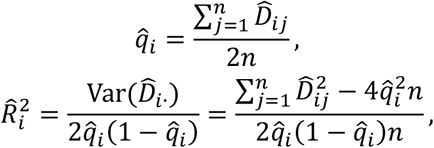

where *n* is the sample size and Var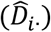 is the variance of 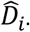 across samples. We calculate SNP-level missing rate as the proportion of genotypes being missing or with 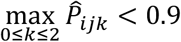. To ensure the data quality, we exclude SNPs with MAF < 0.01, HWE *p*-value < 10^−6^, 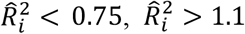, or missing rate > 0.1.

### Simulation data

To examine the performance of our genotype calling algorithm, we conducted simulations with two parameter settings, mimicking the RAI distributions of Type I probes (*α*_O_ = 3, *β*_O_ = 35, *α*_1_ = *β*_1_ = 20, *α*_2_ = 65, *β*_2_ = 4, *λ* = 0.025) and Type II probes (*α*_O_ = 2, *β*_O_ = 20, *α*_1_ = 35, *β*_1_ = 40, *α*_2_ = 40, *β*_2_ = 3, *λ* = 0.015), respectively. Let *m* and *n* be the number of probes and samples. We chose four values of *m* (50, 100, 200, and 400 for Type I probes; 500, 1,000, 2,000, and 4,000 for Type II probes) and six values of *n* (100, 200, 400, 800, 1,600, and 3,200) in different simulations. In each simulation, we first randomly drew allele frequencies of *m* common SNPs (MAF > 0.05) from the 1KGP data, denoted as *q*_*i*_ for SNP *i*. We then simulated an *m*×*n* genotype matrix (***T***) by drawing genotypes of SNP *i* from a binomial distribution with probability *q*_*i*_. Next, we randomly set a small fraction (*λ*) of genotypes in ***T*** to missing. Finally, we simulated an *m*×*n* RAI matrix (***X***) by drawing *X*_*ij*_ from U(0,1) if *T*_*ij*_ is missing and from Beta(*α*_*k*_, *β*_*k*_) if *T*_*ij*_ = *k*, where *k* = 0, 1, or 2. In total, we generated 48 (=2×4×6) sets of simulations based on combinations of parameter settings, sample sizes, and numbers of probes. For each simulation, we repeated 20 times to evaluate the mean and standard error (SE) of the accuracy of our methods in estimating the parameters and genotypes.

### Inference of population structure

We inferred population structure for the AIBL samples using the LASER method [23,40]. Briefly, we first defined an ancestral space using the top four principal components (PCs) of principal component analysis (PCA) of the 1KGP samples, because the top four PCs of 1KGP could separate major continental groups [34]. We then used the *trace* program in LASER to project each study sample into the ancestral space based on genotypes from MethylGenotyper.

### Estimation of kinship coefficient

For DNAm data, we estimated kinship coefficients by SEEKIN [6], which accounted for the uncertainty in the inferred genotypes through the dosage *R*2 for each SNP. We used the SEEKIN-hom and SEEKIN-het estimators for samples from the DFTJ and AIBL cohorts, respectively. The SEEKIN-het estimator can account for the diverse ancestry by introducing individual-specific allele frequencies [6]. Briefly, given an ancestral space defined by the top four PCs of the 1KGP samples, we modeled genotypes with a linear function of the top four PCs:

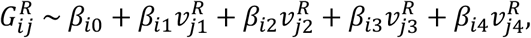

where 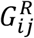 indicates the genotype at SNP *i* of individual *j* from the reference samples, 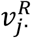 indicates the PC coordinates of individual *j*, and *β*_*i*_ are the regression coefficients for SNP *i*. Denoting 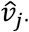 as the projected PC coordinates of the *j-*th study sample, the individual-specific AF 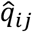 can be estimated as

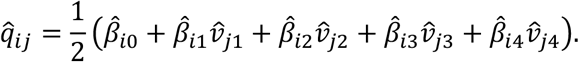

We truncated 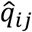 at 0.001 and 0.999 for values outside the boundary.

For comparison, we estimated kinship coefficients using GWAS data of the DFTJ and AIBL cohorts. For the DFTJ cohort, we applied SEEKIN-hom [6] on GWAS data of 286,727 SNPs with MAF > 0.01 and linkage disequilibrium (LD) *r*^2^ < 0.5. For the AIBL cohort, we applied PC-Relate [10] on GWAS data of 113,690 SNPs with MAF > 0.05 and LD *r*^2^ < 0.2. Based on kinship coefficients estimated from GWAS data, we classified relatedness into duplicates, 1^st^ degree, 2^nd^ degree, 3^rd^ degree, and unrelated pairs according to the cutoffs described in Manichaikul *et al*. [8]. In addition, we grouped 2^nd^ degree or closer relatedness (kinship coefficient > 2^-3.5^) as the positive set and the rest as the negative set to compute the following statistics for DNAm-based kinship classification:

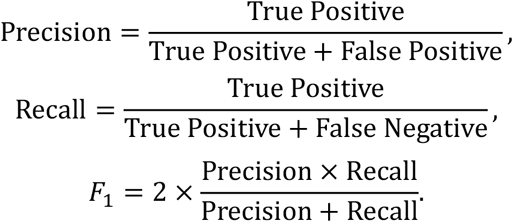

## Data availability

The DFTJ data are available through reasonable request to the corresponding authors, governed by the Regulations on the Management of Human Genetic Resources in China. The AIBL DNAm data are available at https://www.ncbi.nlm.nih.gov/geo/query/acc.cgi?acc=GSE153712. The AIBL genetic data are accessible to all interested parties through an Expression of Interest procedure governed by the AIBL Data Use Agreement (https://aibl.csiro.au/awd).

## Code availability

The R package of MethylGenotyper is publicly available at https://github.com/Yi-Jiang/MethylGenotyper.

## Acknowledgements

This study is supported by grants from National Natural Science Foundation of China (82325044, 82021005), the China Postdoctoral Science Foundation (2021M701318), the Natural Science Fund for Distinguished Young Scholars of Hubei Province (2022CFA046), and the Fundamental Research Funds for the Central Universities (HUST: 2019kfyXJJS036, 2023BR030). Genetic and DNAm data in the AIBL study were funded by the National Health and Medical Research Council (GNT1161706, GNT1151854). We thank all the investigators of the DFTJ cohort and the AIBL study for their contributions.

## Author contributions

CW and SC conceived and supervised the study, generated data, and interpreted the results. YJ and MQ analyzed the data, interpreted the results, and drafted the paper. YJ implemented the R package. MJ and XJ analyzed the DFTJ GWAS data. HG contributed to the generation and analysis of the DFTJ data. SF, TP, SML and CLM were responsible for generating, collating, and analyzing the AIBL data. All authors revised and approved the paper.

