## Supplemental figures for "Accurate estimation of SNP genotypes and genetic relatedness from DNA methylation data"

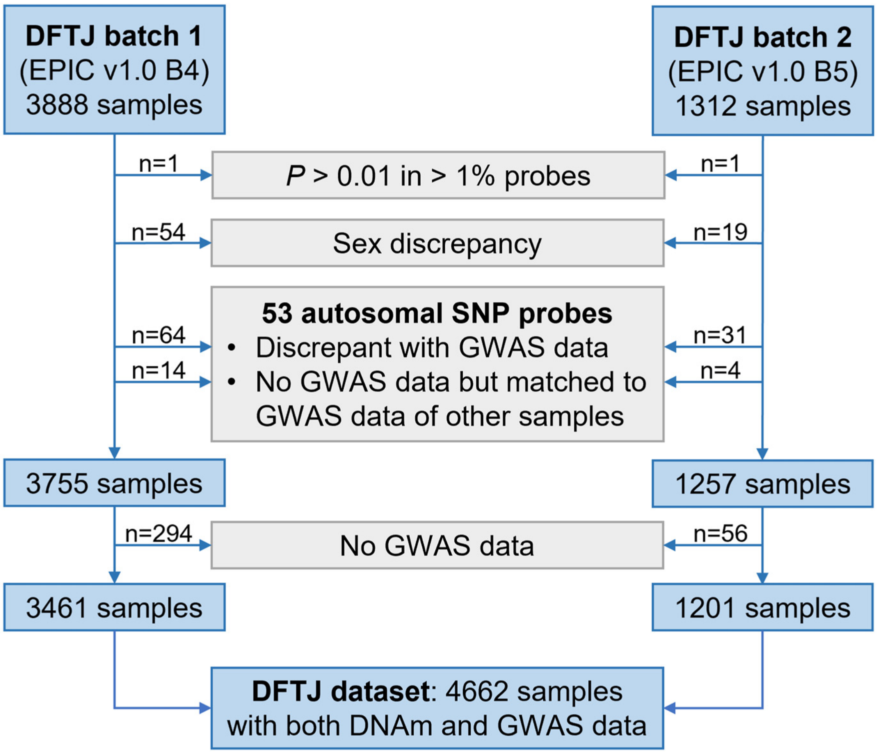


**Figure S1. Sample filtering procedure of the DFTJ dataset.** We used ewastools to infer sexes and genotypes at SNP probes from DNAm data. We excluded samples with inferred sex different from self-reported sex, with discordant genotypes at >10 SNP probes compared to his/her own GWAS data, or with discordant genotypes at ≤3 SNPs probes compared to GWAS data of other samples. The final 4,662 samples were from 4,542 individuals, including 114 individuals with duplicate measurements and 3 individuals with triplicate measurements.

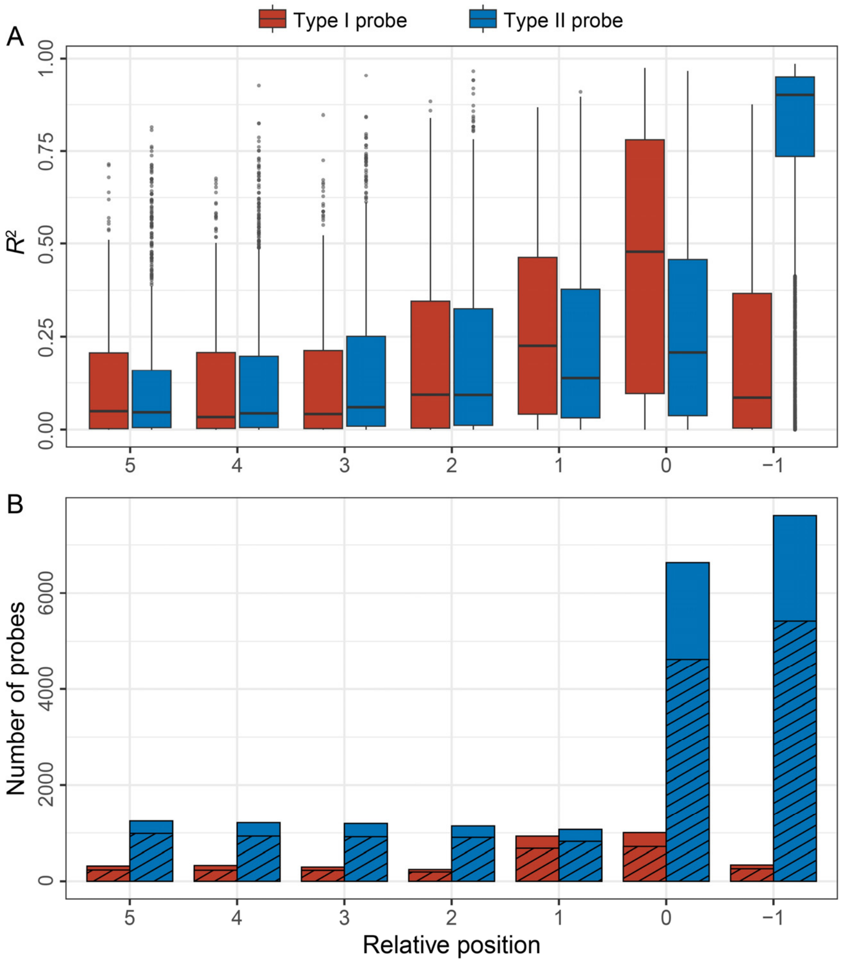


**Figure S2. Correlation between DNAm intensity and genotypes of nearby SNPs in the DFTJ dataset. (A)** Squared correlation ($R^2$) between DNAm β-values (after regressing out age, sex, BMI, smoking status, sample plates and six immune cell type proportions) and genotypes of nearby SNP as a function of their physical distance to the 3' end of the probes. The extension base was marked as -1. **(B)** Number of probes with SNP at different positions relative to the 3' end of the probes. Shaded areas indicate the number of SNPs with genotypes available in the DFTJ GWAS data (*i.e.*, SNPs used to calculate $R^2$ in panel A).

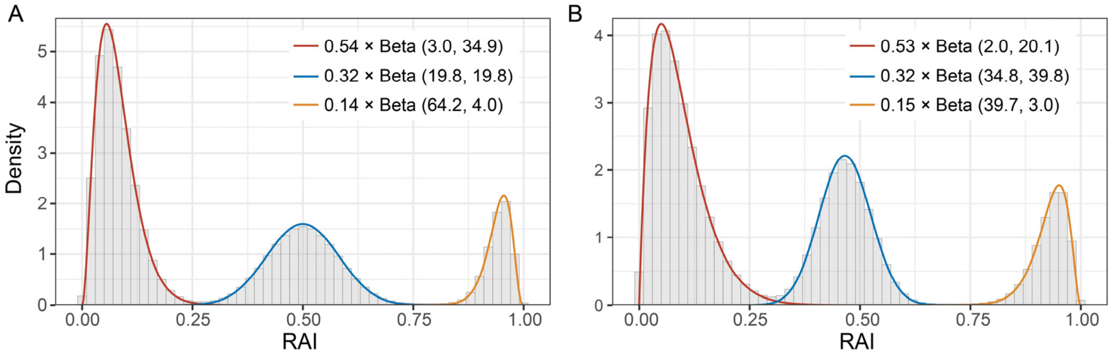


**Figure S3. Distribution of RAIs in simulation data. (A)** Histograms of RAIs across all samples in simulation data matching Type I probe features (true parameters: $\alpha_0 = 3, \beta_0 = 35, \alpha_1 = \beta_1 = 20, \alpha_2 = 65, \beta_2 = 4$, and $\lambda = 0.025$). The simulation data include 400 SNPs and 3,200 samples. **(B)** Histograms of RAIs across all samples in simulation data matching Type II probe features (true parameters: $\alpha_0 = 2, \beta_0 = 20, \alpha_1 = 35, \beta_1 = 40, \alpha_2 = 40, \beta_2 = 3$, and $\lambda = 0.015$). The simulation data include 4,000 SNPs and 3,200 samples. The smooth lines show mixtures of three beta distributions.

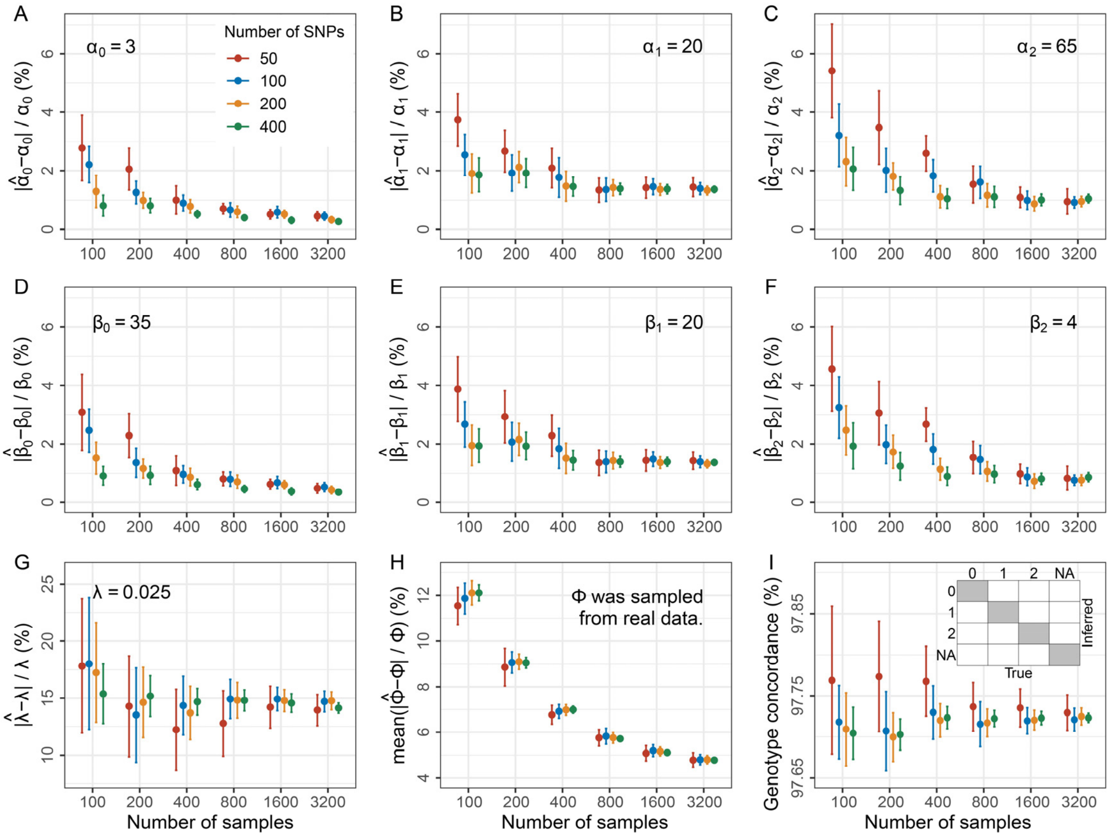


**Figure S4. Performance of MethylGenotyper on simulation data mimicking the RAI distribution of Type I probes.** The simulation data was produced by specifying the parameters for the mixed three beta distributions ($\boldsymbol{\alpha}$ and $\boldsymbol{\beta}$) and one uniform distribution ($\lambda$), based on the characteristics of Type I probes in DFTJ. AFs ($\boldsymbol{\phi}$) randomly sampled from 1KGP were used to infer the mix proportions. **(A-G)** Error rates of the estimated parameters. **(A-C)** $\alpha$ of genotype 0, 1, and 2. **(D-F)** $\beta$ of genotype 0, 1, and 2. **(G)** The outlier probability $\lambda$. **(H)** Mean error rates of $\boldsymbol{\phi}$. **(I)** Genotype concordances, which were calculated by dividing the number of genotypes in the shade areas by the total number of genotypes. The simulation procedure was repeated 20 times to obtain the means and corresponding 95% CIs (mean ± 1.96×standard errors).

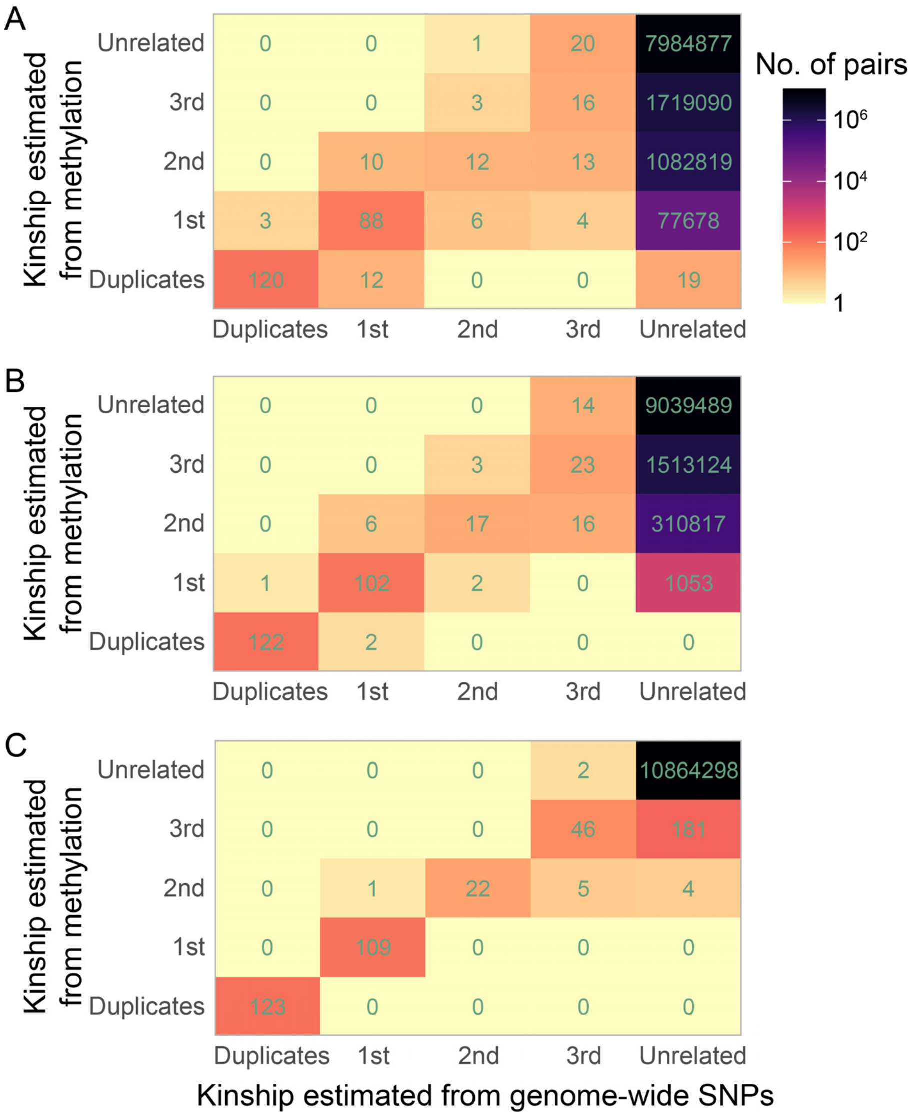


**Figure S5. Comparison of kinship estimated from methylation and from genome-wide SNPs in the DFTJ dataset.** The x-axis indicates kinship estimated from 286,727 genome-wide SNPs (gold standard). The y-axis indicates kinship estimated from methylation data based on **(A)** 53 SNP probes, **(B)** 53 SNP probes and 111 Type I probes, and **(C)** 53 SNP probes, 111 Type I probes, and 4,155 Type II probes.

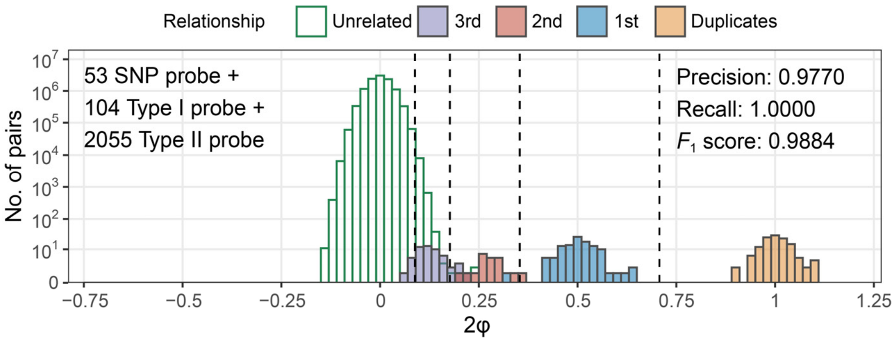


**Figure S6. Performance of kinship estimation based on MethylGenotyper-inferred genotypes for probes in 450K in the DFTJ dataset.** We calculated the kinship coefficient φ based on 2,212 probes in 450K. Only probes with dosage $R^2 > 0.75$ were used. Colored bars represent different relationship types determined by the gold standard. Dashed lines are the inference criteria of ϕ. Precision, recall, and $F_1$ score were calculated by comparing with the gold standard, with sample pairs of 2nd degree or closer being taken as positive.

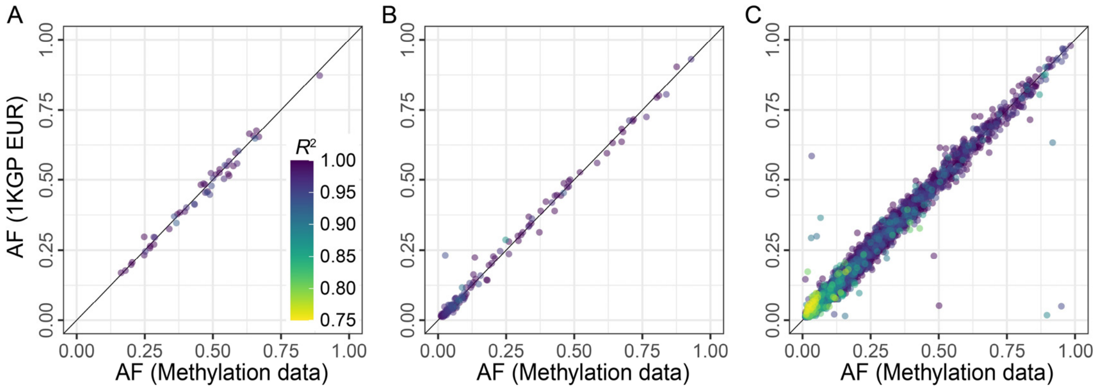


**Figure S7. Comparison of AFs from the AIBL methylation data and the 1KGP European genotyping data.** **(A)** SNP probes, **(B)** Type I probes, **(C)** Type II probes. Genotypes from methylation data were called from MethylGenotyper. Each point represents a SNP, colored by the estimated dosage $R^2$. Only SNPs passed QC were shown.
